# Interaction modifications lead to greater robustness than pairwise non-trophic effects in food webs

**DOI:** 10.1101/377473

**Authors:** J. Christopher D. Terry, Rebecca J. Morris, Michael B. Bonsall

## Abstract

1. Considerable emphasis has been placed recently on the importance of incorporating non-trophic effects in to our understanding of ecological networks. Interaction modifications are well established as generating strong non-trophic impacts by modulating the strength of inter-specific interactions.
2. For simplicity and comparison with direct interactions within a network context, the consequences of interaction modifications have often been described as direct pairwise interactions. The consequences of this assumption have not been examined in non-equilibrium settings where unexpected consequences of interaction modifications are most likely.
3. To test the distinct dynamic nature of these ‘higher-order’ effects we directly compare, using dynamic simulations, the robustness to extinctions under perturbation of systems where interaction modifications are either explicitly modelled or represented by corresponding equivalent pairwise non-trophic interactions.
4. Full, multi-species representations of interaction modifications resulted in a greater robustness to extinctions compared to equivalent pairwise effects. Explanations for this increased stability despite apparent greater dynamic complexity can be found in additional routes for dynamic feedbacks. Furthermore, interaction modifications changed the relative vulnerability of species to extinction from those trophically connected close to the perturbed species towards those receiving a large number of modifications.
5. Future empirical and theoretical research into non-trophic effects should distinguish interaction modifications from direct pairwise effects in order to maximise information about the system dynamics. Interaction modifications have the potential to shift expectations of species vulnerability based exclusively on trophic networks.

## Introduction

There is a building appreciation that to improve our understanding of the population dynamics of ecological communities it is necessary to move beyond studies that focus on a single interaction process at a time (Kéfi et al., 2012; Levine, Bascompte, Adler, & Allesina, 2017). Trophic interaction modifications (Terry, Morris, & Bonsall, 2017; Wootton, 1993) occur when a consumer-resource interaction is modulated by additional species. They are a class of higher-order processes since their effects are not fundamentally pairwise. Examples include associational defences (P.Barbosa et al., 2009), fear effects (Sih, Englund, & Wooster, 1998), certain impacts of ecosystem engineers (Sanders et al., 2014) and foraging choices (Abrams, 2010). It has been empirically demonstrated that many strong non-trophic effects are caused by such processes (Ohgushi, Schmitz, & Holt, 2012; Preisser, Bolnick, & Benard, 2005; Werner & Peacor, 2003) with large implications for community structure and dynamics (M. Barbosa, Fernandes, Lewis, & Morris, 2017; Donohue et al., 2017; Matassa, Ewanchuk, & Trussell, 2018; van Veen, van Holland, & Godfray, 2005).

Approaches to understanding interaction modifications often try to distil the inherently multi-species process into a pairwise non-trophic effect (or ‘trait-mediated indirect effect’) from the modifier species onto one or both of the recipient species (Okuyama & Bolker, 2007). This allows the direct comparison of non-trophic and trophic interaction strengths (Preisser et al., 2005) and network structure (Pilosof, Porter, Pascual, & Kéfi, 2017) but is a representation of a different class of dynamic process (Terry et al., 2017). While this simplification is fully justified in studies of communities at equilibrium (Grilli, Barabás, Michalska-Smith, & Allesina, 2017), the consequences of this assumption in a transient, fluctuating or heavily perturbed system has yet to be fully explored. Previous studies introducing interaction modifications to trophic networks (Arditi, Michalski, & Hirzel, 2005; Goudard & Loreau, 2008; Lin & Sutherland, 2014) have demonstrated their potential impact on the dynamics of ecosystems, but not whether this is attributable to the higher-order nature of interaction modifications, or rather shifts in connectance and interaction strength.

One such important case is the dynamics of ecological systems in the face of species removal, where there is the potential for secondary extinctions and eventually the collapse of the ecosystem (Dunne, Williams, & Martinez, 2002). This aspect of stability, often described as ‘robustness’, is an important question both from the perspective of managing anthropogenic change and in terms of understanding the fundamental stability of ecological communities. Since empirically testing how whole communities respond to extinctions can be difficult or impossible (although see Sanders et al. (2018)), a number of studies have attempted to determine the properties that make ecological communities robust through simulation (Dunne & Williams, 2009; Säterberg, Sellman, & Ebenman, 2013). However, incorporating the acknowledged flexibility of ecological networks is a perennial challenge for such studies (Montoya, Pimm, & Solé, 2006).

The impact on the robustness of ecological networks of one specific subset of interaction modifications, those caused by the flexible foraging in response to resource availability, has been examined by a number of studies (Valdovinos, Ramos-Jiliberto, Garay-Narváez, Urbani, & Dunne, 2010). Approaches have included topological rewiring (Gilljam, Curtsdotter, & Ebenman, 2015; Kaiser-Bunbury, Muff, Memmott, Müller, & Caflisch, 2010; Staniczenko, Lewis, Jones, & Reed-Tsochas, 2010; Thierry et al., 2011), multi-species functional responses (Uchida & Drossel, 2007) and adaptive foraging models (Kondoh, 2003). These models showed that the additional dynamic process impacted robustness in contrasting directions, but only addressed a restricted subset of interaction modifications caused by predator switching. However, consumption rates are influenced by more than the just the choice of prey available to them. Interaction modifications can also be caused by the threats to the consumer (Suraci, Clinchy, Dill, Roberts, & Zanette, 2016), facilitation (Bruno, Stachowicz, & Bertness, 2003), associational susceptibility (Underwood, Inouye, & Hambäck, 2014) or mutualistic defence (Holland, Ness, Boyle, & Bronstein, 2005), amongst others (Ohgushi et al., 2012). This introduces a considerable number of additional links between species to ecological communities, yet studies of generic interaction modifications in large networks are limited (Arditi et al., 2005; Bairey, Kelsic, & Kishony, 2016; Garay-Narváez & Ramos-Jiliberto, 2009; Goudard & Loreau, 2008; Lin & Sutherland, 2014), with most theoretical analyses of interaction modifications focussing on small community units (Abrams, 2010; Bolker, Holyoak, Křivan, Rowe, & Schmitz, 2003; Holt & Barfield, 2013).

Calls to incorporate the full panoply of non-trophic effects into our understanding of ecological networks have built substantially in recent years (Fontaine et al., 2011; Ings et al., 2009; Kéfi et al., 2012; Levine et al., 2017; Ohgushi et al., 2012; Olff et al., 2009) and the first empirical inventories are being established (Kéfi et al., 2015; Kéfi, Miele, Wieters, Navarrete, & Berlow, 2016). Theoretical analyses can play a significant role in motivating the empirical construction of networks, identifying the information necessary to best understand these systems. Here we demonstrate the distinctive nature of interaction modifications compared to pairwise non-trophic effects through a direct standardised comparison of their impacts on the robustness of large artificial networks. We then examine the role of interaction modifications in determining the consequences of species loss and species level vulnerability to secondary extinction.

## Methods

We conducted our analyses using model food webs where system dynamics are derived from metabolic scaling relationships. As detailed below, interaction modifications were introduced to communities at an initial equilibrium before mortality was added to a single species at a time and examined the resultant extinctions after the model was integrated to a new equilibrium.

### Bio-energetic Model

The change in biomass density, *B_i_*, of each species in the community was modelled using a simple Lotka-Volterra type model with a linear (Holling Type I) functional response and logistic intrinsic growth rates, parameterised using body-mass relationships:

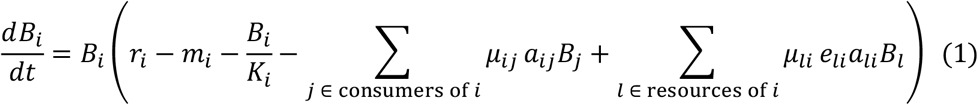

Each species was assigned a body mass (*M_i_*) drawn from a distribution based on their trophic level (see SI 1) which was then used to calculate further parameters using quarter-power body-mass scaling laws (Yodzis & Innes, 1992). Relative intrinsic growth or metabolic loss rates, *r_i_*, were set at 1 for all producers (trophic level 1) and *r_i_*= −0.1*M*^−0.25^ for each consumer (trophic level ≥2). Consumer-specific consumption rates were set at *a_ij_* =*w_j_M*^−0.25^, where the generality term *ω_j_* was set to 1/*n*, the number of resources of each consumer *j*. Assimilation efficiencies, *e_ij_*, for each trophic interaction were drawn from a uniform distribution, *e_ij_* ~ 𝒰 (0.05: 0.15). Carrying capacities, *K_i_*, were drawn from *K_i_*~𝒰 (1: 10) for producers (to introduce a moderate degree of self-regulation) and *K_i_*~10 𝒰 (2:3) for consumers (considerably higher than the starting populations and so introducing only a small amount of self-regulation). Initially, external mortality, *m_i_*, was set to 0 and modification terms, *μ_ij_*, set to 1.

Trophic topology and population densities of starting communities were determined as follows. Initial trophic topologies were generated using the niche model (Williams & Martinez, 2000), with 35 species and a connectance of 0.14, removing cannibalistic interactions. Each population density was initially set to 10 and these systems were then numerically integrated to a stable equilibrium (see SI 4 for criteria). This was repeated until 200 fully-connected communities with at least 18 persisting species were generated to be used as starting communities for the robustness tests. Properties of the starting communities are described in SI 2.

### Specification of TIMs

Trophic interaction modifications were introduced to these communities through modification terms, *μ_ijk_*, that specify the impact of modifier species *k* on the consumption of species *i* by species *j*, with the constraint that *i* ≠ *j* ≠ *k*. These are positive numbers that multiply the attack rate as a function (detailed in the next section) of the divergence of the biomass density of a modifying species, *B_k_*, from its starting equilibrium value 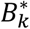. Where there are multiple modifications of the same interaction these were assumed to combine multiplicatively *μ_ij_* = ∏^*k*^ *μ_ijk_* (Golubski & Abrams, 2011; Goudard & Loreau, 2008).

When *μ_ijk_* < 1, the interaction is weakened and when *μ_ijk_* > 1 the interaction is strengthened. In our model *μ_ijk_* can not be negative, preventing the reversal of the direction of the trophic interaction. Where an increase in the modifier species leads to an increase in the strength of the interaction, we describe it as a facilitating TIM. This can be said to be beneficial to the consumer and detrimental to the resource, in the sense of the immediate impact from an increase in the modifier population. We term the reverse situation an interfering TIM. See Figure 1 for diagrammatic explanation of these terms.

**Figure 1.**
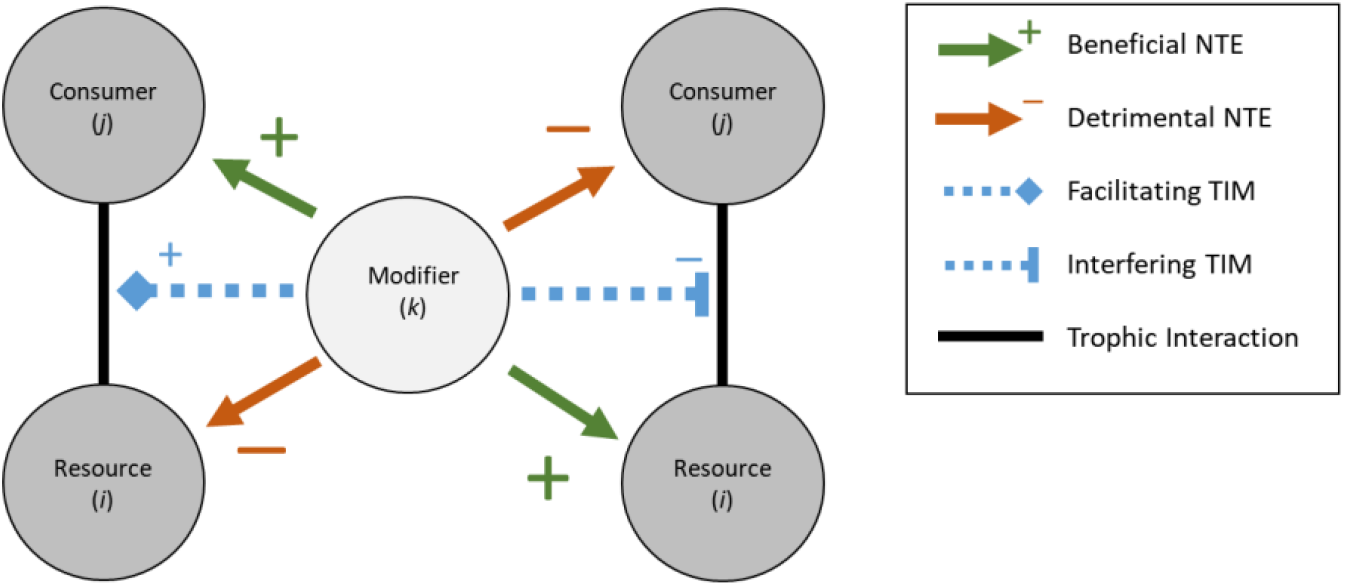
Distinctions between terms used to describe interaction modifications and their impacts in terms of the effect of an increase in the density of the modifier species (centre). ‘Facilitating’ and ‘interfering’ trophic interaction modifications (TIMs, dotted arrows) cause ‘beneficial’ and ‘detrimental’ short term non-trophic effects (NTEs, solid arrows), alternately on the consumer and the resource.

### Functional Form of TIM

To represent the relationship between the density of the modifier species and the modification of the interaction we used a Gompertz sigmoidal curve parameterised to control features of ecological relevance, detailed in full in SI 3. This function links the divergence of *B_k_* from its start point,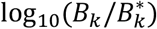, and three control parameters depicted in Figure 2. These are τ (the distance from the starting point the threshold point where the response to a change in modifier density is greatest), α (the direction and maximum rate of change in the modified interaction as the modifier density increases) and σ (the range of magnitudes over which *μ_ijk_* spans). A positive α denotes a facilitatory modification and a negative α an interference modification. A key attribute of our parameterisation is that when 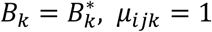. This maintains the original trophic interaction strength when the modifier is at its starting equilibrium point - in effect these strengths are assumed to already incorporate the effect of the modifier to the interaction at the equilibrium.

**Figure 2.**
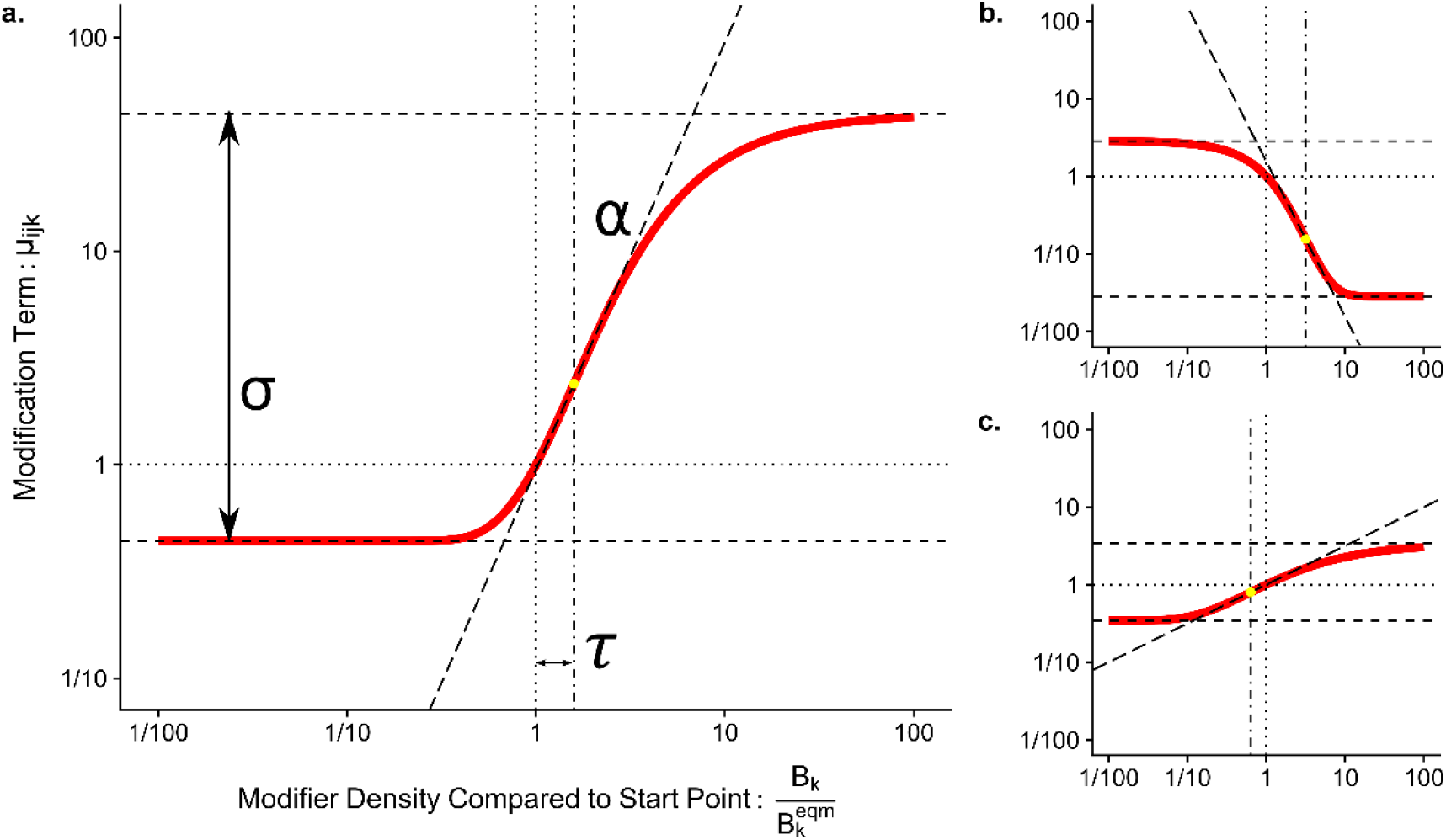
Graphical representation of the relationship between the control parameters and the response of the sigmoidal function used to determine the modification term μ_ijk_ from the density of the modifier B_k_. Panel a) uses a maximum slope (α) of 2, a distance to threshold (τ) of 0.2 and a range of effects (σ) to 2. Note that the function is calculated on a base 10 logarithmic scale. Parameters for panel b): α = -2, τ = 0.5, σ = 2 and panel c): α=0.5, τ = -0.2, σ =1.

### Alternative representation of TIMs as Pairwise Non-Trophic Effects

As an alternative to modelling the impact of trophic interaction modifications in full (using ‘higher order’ terms), equivalent pairwise non-trophic effects (NTEs) can be derived. These match the impact of the full TIM model, the only distinction being the NTE from a modifier *k* to a trophic interactor is no longer dependent on the biomass of the other member of the trophic pair (Figure 3). To maintain parity with the TIM model, this was done by first partitioning the process into trophic and non-trophic components, then fixing the value of the other trophic interactor to the density at the original equilibrium: 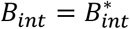. Full steps of the derivation are in SI 6. A trophic interaction onto a resource influenced by a modifier can be partitioned from:

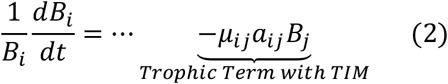

to:

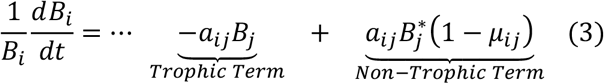

**Figure 3.**
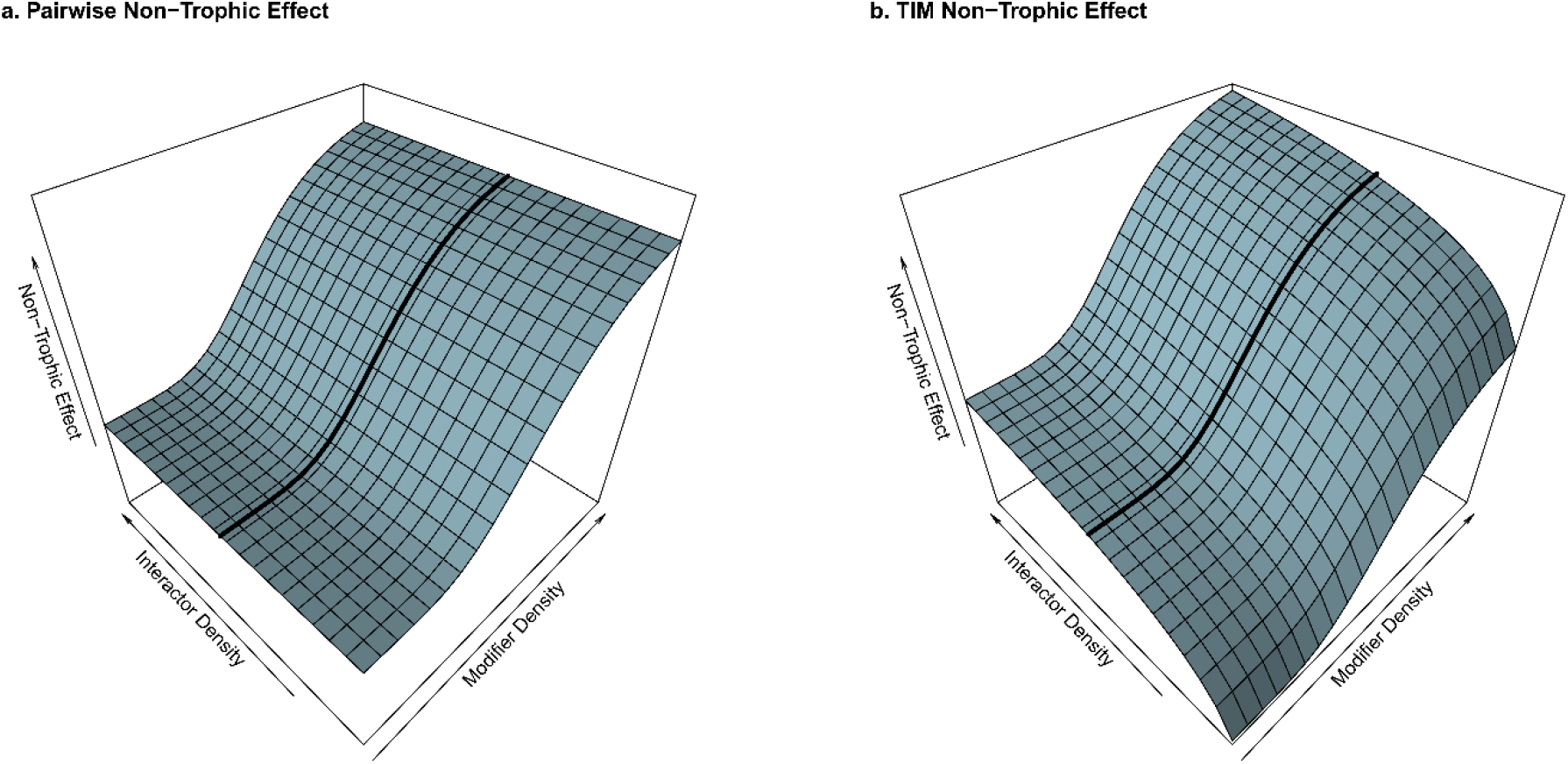
Illustrative response surfaces showing the distinction between non-trophic effects caused by pairwise interactions and those from a higher-order interaction modification process involving three species. Per-capita effects are shown in the vertical axis on a logarithmic scale. In both cases, the response to the modifier is identical at the starting density of the trophic interaction partner of the focal species, shown by the black line. However, the density of the trophic interaction partner of the focal species (left-hand axis) does not affect the non-trophic effect in the pairwise case (a), but does with the full interaction modification model (b). Note also that as the interactor’s density becomes small, the strength of the non-trophic effect rapidly declines.

The corresponding terms for the interactions onto the consumer, *j*, are:

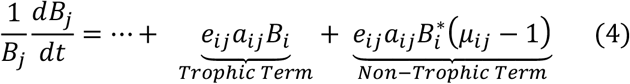

As an example, a facilitating TIM, when *B_k_* increased, would lead to a *μ_ij_* greater than 1. This would lead to a negative NTE on the resource *i* (independent of *B_j_*) and a positive NTE on the consumer *j* (independent of *B_i_*). This partitioning process is only straightforward when each trophic interaction is modified by at most one modifier species because of the synergistic relationship between multiple modifiers assumed by our model.

### Test 1. Comparison of robustness between pairwise NTE and higher-order TIMs

To compare the consequences of introducing TIMs by the two approaches, we conducted three sets of robustness tests using the same set of trophic networks. The first using the full-TIM model, the second using the TIMs that had been converted to pairwise form, and a third case without any TIMs.

We randomly added TIMs to the set of initial communities such that each potential modification (combination of trophic interaction and third species) had an equal 0.05 chance of existing. For this test each interaction was modified by at most one other species to allow the conversion to non-trophic effects. Shape parameters for each TIM were drawn from uniform distributions: slope α~ 𝒰(−4: 4), range σ ~ 𝒰 (0.1: 4) and threshold τ ~ 𝒰 (−1: 1). The location of TIMs and their shape parameters were identical between the full-TIM and pairwise TIM cases.

For each robustness test, the external mortality rate (*m_i_*) of a single species (to which we will refer for convenience as the ‘targeted’ species, although the mortality could be attributable to a range of non-directed processes) was then set to 1, the system numerically integrated to a new steady state and the status of each species in the resultant community assessed. Species were considered ‘extinct’ if their biomass fell permanently below 10^−4^ (several orders of magnitude below the starting values of most of the populations, SI 2), ‘functionally extinct’ if their biomass fell to below 1/10^th^ of their starting value and considered to have ‘exploded’ if the final density was over 10 times the starting value. Robustness tests were repeated targeting each species in turn, for each community, to give a total of 3736 completed robustness tests (93.8% successful integrations, SI 4). All analyses were carried out in R v.3.5.0 (R Core Team, 2018) using the *deSolve* numerical integration package (Soetaert, Petzoldt, & Setzer, 2010).

### Test 2. TIMs and distribution of extinctions

To examine the relationship between the distribution of TIMs and consequent secondary extinctions, the robustness tests of the full TIM case as described above was repeated with a higher occurrence rate of TIMs (0.08) and a relaxation of the assumption that only one modification can affect each trophic interaction. Results are reported for parameters drawn from α ~ 𝒰 (−3: 3), σ~ 𝒰 (0.1: 3), τ~ 𝒰 (−1: 1). Further tests, with different TIM occurrence rates and distributions of shape parameters, reached qualitatively similar results. Properties of the community and the relationship between the ‘targeted’ species and the extinct species were then calculated.

Firstly, for each robustness test, the trophic distance from the targeted species to each secondarily extinct species was calculated. This is the number of trophic links between the targeted species and the secondarily extinct species by the shortest route using the starting network. For this only the trophic interactions were considered, not the TIMs. To generate a baseline to compare to, the trophic distance from the targeted species to every other species was calculated.

Secondly, the NTEs affecting each extinct species was counted and classified by whether an increase in the modifier would be beneficial or detrimental for the focal species. This was done for all NTEs affecting the extinct species, and separately for just those derived from TIMs where the targeted species was the modifier species. In our model, species that are involved in a greater number of trophic interactions will tend to be the recipients of a greater number of NTEs. To distinguish the effect of increased trophic degree from the number of NTEs, for each species we calculated the expected number of incoming NTEs based on the number of trophic interactions and the species richness of the network to generate a baseline. This actual number of incoming NTEs (beneficial or detrimental) for each extinct species was then compared to this expectation baseline.

## Results

The introduction of TIMs greatly increased the number of extinctions (Figure 4), but almost half the number of extinctions were observed when the interaction modifications were modelled directly compared to when they were modelled as pairwise effects (mean difference = 5.4, t-test paired by community and targeted species ID: p<0.0001, t = 61.5, df = 3941). The number of functional extinctions was very similar to full extinctions (means: No-TIM: 1.98, Pairwise: 11.33, Full-TIM: 5.85). Population explosions were rare in all scenarios. Without NTEs there was a mean of 0.09 explosions per robustness test, with nearly twice as many under pairwise NTE (0.189). The TIM case had the fewest, only 0.05 explosions per robustness test.

**Figure 4.**
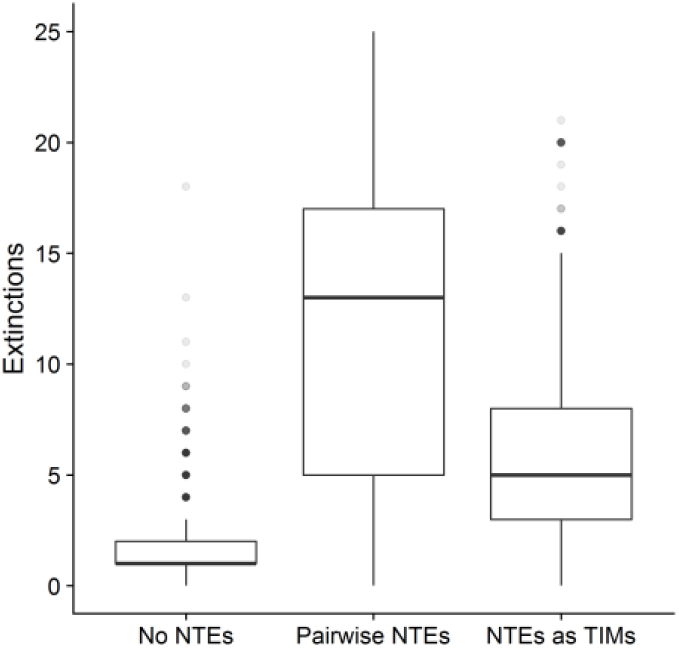
Boxplot showing that robustness was significantly lower when TIMs were modelled as pairwise interactions rather than directly as interaction modifications. Both cases produced significantly more extinctions than the no-TIM case.

In robustness tests without TIMs nearly 60% of secondary extinctions were species directly trophically linked to the targeted species (Figure 5). The introduction of TIMs shifted the trophic distance (the number of trophic links between species by the shortest route) much closer to the underlying distribution of trophic path lengths (Figure 5).

**Figure 5.**
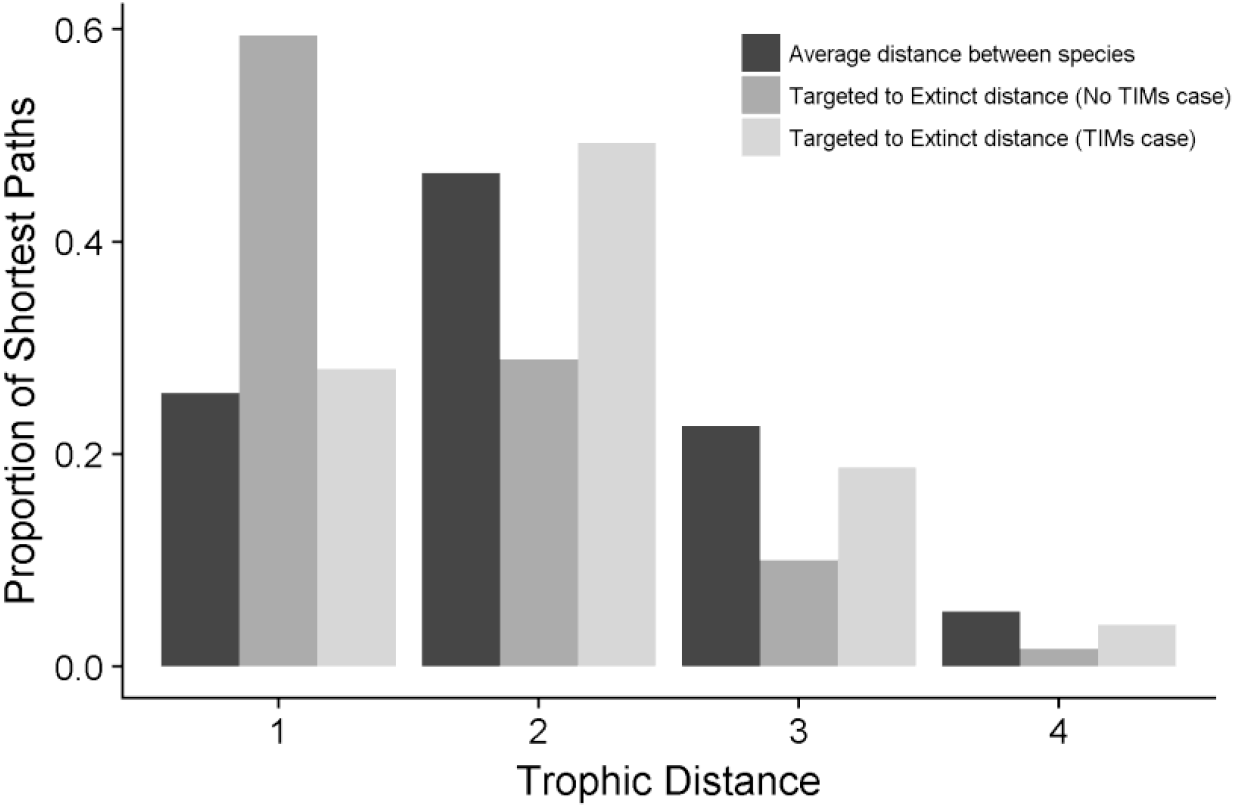
Distribution of trophic distances between all species in the communities (black) and between targeted species and secondarily extinct species in communities without TIMs (dark grey) and communities with TIMs (light grey). The distance from the targeted species to the secondarily extinct species in the TIM case was intermediate between the two baseline cases. Trophic distances above 4 were very rare in our model and not shown.

Secondarily extinct species tended to be recipients of both more beneficial (mean = 4.53) and more detrimental (mean = 4.02) NTEs than would be expected to affect for a random species (mean = 3.79, (t-tests paired by species ID, all p<0.0001, n =3576). When counting just TIMs from the targeted species, secondarily extinct species tended to have more beneficial (mean = 0.289) but fewer detrimental (mean = 0.178) NTEs than the expected number (mean = 0.207,t-tests paired by species ID, all p<0.0001, n =3576).

The targeting of species that caused more TIMs induced more extinctions (Figure 6a, Poisson glm, p<0.0001, n= 3825), with each additional TIM emanating from the targeting species leading on average to an additional 7% secondary extinctions. Model fits splitting the sign of the TIMs (Figure 6b) indicated that each additional interference TIM caused a greater increase in the number of extinctions (+10.4%), compared to each additional facilitatory TIM (+4.1%). Including the trophic connectance of the web as an additional predictor variable did not give a significant improvement in the model fit (*χ*^2^ comparison of models, p=0.113).

**Figure 6.**
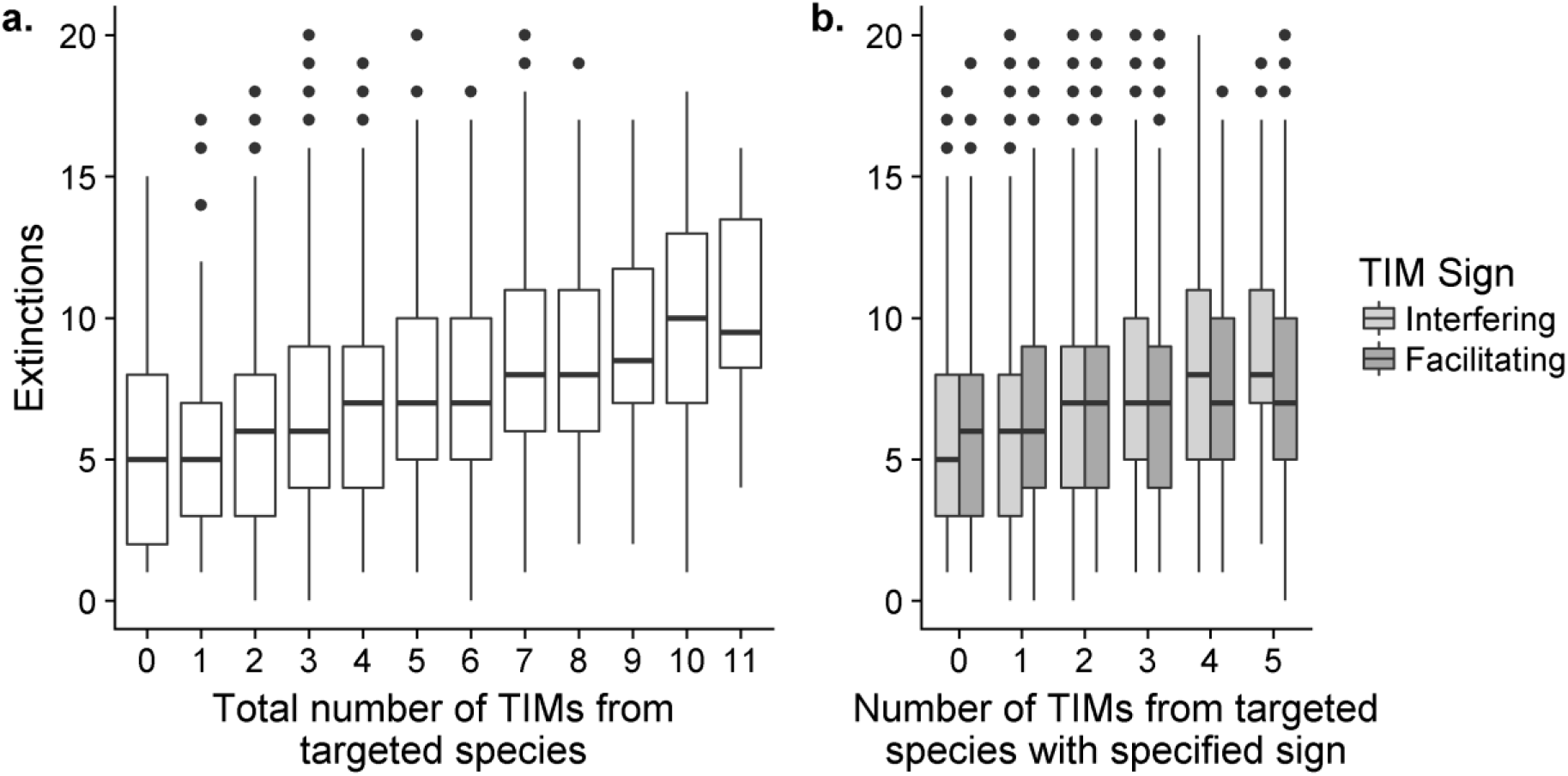
Boxplots showing the increase in the resultant number of extinctions as the number of TIMs caused by the targeted species increased. Panel a) shows the response to the total number of TIMs, panel b) shows the distinct responses to the number of each sign of TIM.

## Discussion

Here we have examined the ecological robustness of networks to demonstrate that interaction modifications have distinct dynamic effects. In ecological systems, efforts to include non-trophic effects into food webs (Fontaine et al., 2011; Pilosof et al., 2017) should take into account the higher-order nature of interaction modifications. Without quantitative information of the topological and strength distribution of TIMs it is not yet possible to precisely calculate their impacts relative to trophic interactions. Nevertheless, it is clear that they have the potential to change our expectation of both the relative vulnerability of species to extinction and the impact the loss of certain species will have (Donohue et al., 2017).

We found that despite an increase in the number of ‘moving parts’ the full TIM model was more robust than the exclusively pairwise model. Generally, unstructured dynamic models of community stability suggest that a higher complexity is destabilising (May, 1972; Uchida & Drossel, 2007) making our result somewhat surprising, but there are a number of explanations. The overall number of direct relationships between species through all types of interaction (dynamic connectance) between our two representations of TIMs is the same. In part, our result can be attributed to the tendency for an overall decline in population densities under the perturbation. In the full-TIM model the strength (and disruptive influence) of the impact is dependent on the product of two species and so (in the context of generally declining species densities) will decline faster than when the strength is dependent on just one species. Weaker effects between species due to TIMs will reduce the potential for cascading effects. Furthermore, if the density of either a consumer or resource falls to zero, any TIM impact on the species it is trophically liked to would will also fall to zero under the full TIM model. However, with the pairwise model the NTE is maintained even though the mechanism is no longer extant. Hence, as species become extinct the effective dynamic connectance of the full TIM model declines faster than in the pairwise case.

There are also differences between the full-TIM and the pairwise models in terms of the prevalence and rapidity of local feedback loops. Feedbacks modulating the impact of TIMs are heavily dependent on the relative speed and strength of multiple feedback loops, which in turn are derived from properties of individual species. Despite our relatively simple model these loops are challenging to trace in large complex systems (Dambacher & Ramos-Jiliberto, 2007), limiting qualitative analysis. Nonetheless, as the impact of the more complex full-TIM model is dependent on more species, in general, it appears that mitigating feedback loops can act faster in the full-TIM case. For example, the effects caused by an interaction modification (however modelled) will cause an immediate increase in one affected species and decrease in the other. In the full-TIM model, where the strength of the interaction modification is dependent on *B_i_*× *B_j_*, at least for a short time the strength of the impact of the TIM on the consumer and the resource will be stay relatively constant since *B_i_* and *B_j_* move in opposite directions. In the pairwise model the strength of the NTE on each species will rapidly diverge with the potential to generate unrestricted positive effects on one of the populations, with disruptive effects for the rest of the community. Over longer time-scales changes in consumption rate lead to a consistent changes in the equilibrium values of both consumer and resource, as can be demonstrated algebraically. In a simple module, a heightened consumption rate leads to greater suppression of the resource which in turn can support a reduced population of consumers (e.g. Morin, 2011). These distinct phases of responses to a change in consumption rate, immediate biomass shifts in opposite directions before eventually moving towards a consistent direction, highlights the multiple timescales over which TIMs operate. The analysis of the contributing factors to these feedback loops would be a profitable area for future work.

At least in this simple model, it appears that TIMs determine extinction risk in a somewhat predictable way. The extinction risk is shifted from those species that are closely trophically connected to perturbed species, to be more evenly distributed throughout the network. Overall, the species that went extinct tended to be those that were affected by more TIMs. When considering NTEs from all species, this was true of both ‘beneficial’ and ‘detrimental’ NTEs. Those species that directly benefit from the presence of other species are clearly affected by extinction cascades. However, those species receiving an above-expectation number of detrimental NTEs were also at higher risk of extinction. It would be expected that these species would benefit from the on-average reduction or removal of their inhibitors. Note that the correlation between the increased species level connectance and the number of modifications received was taken into account by comparing expected number of TIMs on a per-species basis.

There are two complementary explanations for the equivalence of detrimental NTEs. Firstly, in a complex model certain species do increase their density, at least for a period of time, which would lead to have negative effect on the focal species. Initial responses to a small reduction in the population of a single species were as commonly positive as negative (51.1% of non-zero growth-rate responses to a 1% density reduction in a random species were positive in the TIM models, SI 5). The low number of species maintaining large increases at the end of the process (what we defined as population explosions) shows that such increases were relatively transient. When considering only TIMs from the targeted species, which always decrease in density, extinct species received fewer than expected detrimental NTEs, supporting this direct explanation. Secondly, in the simple model of population dynamics we used, an increase in the attack rate of a consumer (which would be classed as ‘beneficial’) eventually leads to a reduction in the equilibrium density of the consumer since the population of resource is pushed down to a level that cannot sustain as high a consumer population.

Much previous work has shown that complex networks are stabilised by consistent patterns in key parameters, which can be derived from body-mass scaling rules (Brose, Williams, & Martinez, 2006; Otto, Rall, & Brose, 2007). In our model this source of stability declines, as these patterns are disrupted by interaction modifications that effectively push each *a_ij_* value out of the allometrically specified range in both directions. However, attack rates in real systems do take place in the context of other species and significant disruptions to pairwise interaction strengths derived from laboratory experiments caused by interaction modifications have been empirically observed (Jonsson, Kaartinen, Jonsson, & Bommarco, 2018). Our results suggest that although species whose interactions are aided by other species are indeed more sensitive to extinction, the overall number of relationships with other species is as important and that both potential ‘beneficial’ and ‘detrimental’ processes should be considered on an equal footing.

Our choice of function to represent interaction modifications, although mathematically complex in form, offers certain advantages compared to previous linear (Arditi et al., 2005; Bairey et al., 2016), exponential (Goudard & Loreau, 2008; Lin & Sutherland, 2014) or rational function (Sanders et al., 2014) models. In particular, it has ability to directly and independently control salient features of the function with clear ecological relevance (distance to threshold, maximum rate of change, range between maximum and minimum). The dependence on the relative divergence from an equilibrial starting point rather than the absolute value of the density of the modifier is also likely to be more representative of ecological processes and considerably more straightforward to parameterise in an ecologically meaningful manner.

Nevertheless, this is still a highly simplistic model. The interaction modifications included in this study were introduced at random, in the sense that each potential modification had an equal chance of existing. Considerable stabilising structuring has been observed within non-trophic effects networks (Kéfi et al., 2015; Kéfi, Miele, et al., 2016) but there is not yet sufficient diversity of examples to be able to determine whether there are consistent features across ecosystems. Analyses of the stability of systems at equilibrium demonstrate the key role of the distribution in terms of strength and position relative to the trophic network of interaction modifications (Terry, Bonsall, & Morris, 2018) and the potential for emergent structure from the combination of trophic and non-trophic interaction networks. The distribution of interaction modifications will have important consequences for the use of structural network properties such as trophic coherence, modularity and nestedness that have been demonstrated to affect stability (Johnson, Domínguez-García, Donetti, & Muñoz, 2014; Stouffer & Bascompte, 2011; Thébault & Fontaine, 2010) since total interaction networks, including non-trophic links, may display different patterns.

Interaction modifications do not solely affect consumption rates and there is considerable scope to develop and test theoretical models of additional processes, such as non-trophic interaction modifications and modification of other aspects of consumer functional responses (Kéfi et al., 2012). The assumption of multiplicative combination of multiple TIMs in a synergistic manner in our second analysis is also questionable. While empirical patterns have not been quantified, it is a reasonable estimate that many modification effects act antagonistically to each other (Golubski & Abrams, 2011). For instance, the presence of a second fear-inducing predator may well have less effect than the first, dampening the impact of a change in either modifier population.

Further opportunities are also open to introduce specific accounting for the timescale of changes and the size of perturbations. It is possible that higher-order interactions are stabilising against small perturbations, which may make the system as a whole more susceptible to large impacts, such as the extinction of certain species (Levine et al., 2017). TIMs also have the potential to create the necessary positive feedback structures to maintain alterative stable states (Holt & Barfield, 2013; Kéfi, Holmgren, & Scheffer, 2016). As yet, the prevalence of these features is largely unknown. The speed of interaction modifications themselves can also vary. While many TIMs are behaviourally mediated and occur essentially instantaneously, others are due to accumulated environmental changes (Sanders et al., 2014) or evolutionary processes (Benkman, Siepielski, & Smith, 2012) and operate at somewhat slower timescales.

## Conclusion

In summary, interaction modifications are potent forces that introduce distinct dynamics to ecological networks. This distinctive nature of interaction modifications is of relevance for dynamic systems in many fields that make use of networks (Strogatz, 2001) since our work shows that the complexity of networks is more than the product of connectance and the number of interacting units. Despite long-standing calls for the inclusion of non-trophic effects into the mainstream of ecological network science that has been long dominated by food webs (Ings et al., 2009), and the publication of the first empirical community level non-trophic network (Kéfi et al., 2015), there remains a great number of significant unknowns about the role of non-trophic effects at the network scale. Our work shows that maintaining interaction modifications as distinct processes within empirical and theoretical networks, rather than as pairwise non-trophic effects (Grilli et al., 2017), will enable us to have a more complete understanding of the system dynamics and allow better predictions of community responses to perturbations. This need not require significant additional effort on the part of the original investigator, but would be challenging for others to retroactively discern from published pairwise interaction networks. Analyses of network robustness are used extensively to understand anthropogenic impacts on natural communities (Evans, Pocock, & Memmott, 2013; Kaiser-Bunbury et al., 2017; Säterberg et al., 2013); as the development and analysis of comprehensive interaction networks expands (Kéfi, Miele, et al., 2016), we must incorporate interaction modifications appropriately.

## Acknowledgements

JCDT was funded by the Natural Environmental Research Council through the Oxford Environmental Research Doctoral Training Program (NE/L002612/1).

## Data and Code Availability

All R code used in this study and the generated simulation results are available on the Open Science Framework: DOI: 10.17605/OSF.IO/W83BR

## Author Contributions

JCDT initiated the research, and JCDT, MBB and RJM contributed to the ideas presented in the manuscript. JCDT conducted the research, facilitated by discussions with RJM and MBB. JCDT wrote the manuscript, and all authors contributed to revisions and approved the final version.

